# Sequence assignment validation in protein crystal structure models with checkMySequence

**DOI:** 10.1101/2023.02.17.528951

**Authors:** Grzegorz Chojnowski

## Abstract

Sequence register shifts remain one of the most elusive errors in experimental macromolecular models. They may affect model interpretation and propagate to newly built models from older structures. In a recent publication I have shown that register shifts in cryo-EM models of proteins can be detected using a systematic re-assignment of short model fragments to the target sequence. Here, I show that the same approach can be used to detect register shifts in crystal structure models using standard, model-bias corrected electron-density maps. I describe in detail five register shift errors detected using the method in models deposited in the PDB.

**Synopsis:** I show that *checkMySequence*, an automated method for validating sequence assignment in cryo-EM structures of proteins, can be used for validating crystal structure models.

## 1. Introduction

Macromolecular crystallography (MX), NMR, and more recently cryogenic electron microscopy (cryo-EM) are the methods of choice for the detailed analysis of structures of proteins and their complexes. Over five decades, the efforts of generations of structural biologists using these methods have resulted in over 200,000 macromolecular structures being deposited in the Protein Data Bank (PDB; (Berman *et al*., 2000)), most of which (87%) have been solved by MX. The PDB is an invaluable resource of experimentally determined structures of half of the known protein families (according to InterPro version 92 (Paysan-Lafosse *et al*., 2022)), often in multiple biochemical contexts and conformations, in apo form and with natural or artificial interaction partners. Recently, it has enabled training Artificial Intelligence (AI) tools that extrapolated the available experimentally determined structural information onto virtually any known protein sequence (Jumper *et al*., 2021, Baek *et al*., 2021).

Despite the unquestionable value of the accumulated knowledge, the PDB is also known to contain many models that are not fully correct. The issue of error propagation from PDB models to AI-based methods remains an open question (Jones & Thornton, 2022). Currently, it seems that AI-predicted models are an excellent aid in building and correcting experimental structures (Terwilliger *et al*., 2022). However, the structures that could have been validated experimentally constitute only a tiny fraction of almost 200 million predicted models available already in the AlphaFold2 database (Varadi *et al*., 2022). Therefore, the importance of extensive validation of both newly determined experimental models and those already available in the PDB, for which experimental data are available (86% of MX structures), cannot be overemphasised as they provide the most reliable and detailed source of information on macromolecular structures currently available.

The errors in experimental macromolecular models have become easier to detect over time due to the continuous development of model-validation tools. Most importantly, the cross-validation in macromolecular crystallography (splitting reflections into “free” and “work” sets) introduced in the early 1990s helps to avoid gross errors in the models (Brunger, 1992). Local tracing errors can usually be identified as a poor fit between an atomic model and corresponding combined electron density map or prominent difference-density map peaks. Although the maps are calculated using phases derived from a tentative model, which may hinder detection of errors, the refinement programs account for this using sigma-A weighting of map coefficients, which reduces model bias (Read, 1986). The local quality of models is validated using expert systems like PROCHECK (Laskowski *et al*., 1993), WHAT_CHECK (Hooft *et al*., 1996), and MolProbity (Prisant *et al*., 2020) focused on stereochemical plausibility of model coordinates. Multiple map- and geometry-based validation approaches can be also conveniently used e.g., in COOT (Casañal *et al*., 2020) during interactive model building process to identify and correct errors. Finally, a detailed validation precedes the deposition of models to the PDB, which is nowadays an indispensable part of a peer review process in most scientific journals (Gore *et al*., 2017).

Indeed, with the availability of a wide range of model validation techniques, the overall quality of the structural models has improved significantly (Brzezinski *et al*., 2020). It was observed for example that the “*clashscore*” from the MolProbity suite, a sensitive indirect indicator of tracing and map-fit issues, has been steadily improving over time for PDB deposits (Williams *et al*., 2018). At the same time, however, the common usage of Ramachandran plot restraints in refinement often masks model issues making the clear reduction of unusual torsion angles in PDB deposits a weak indicator of model quality improvement (Sobolev *et al*., 2020). This may be confusing to the structural biologists, especially those new to the field. They frequently struggle to distinguish outliers from errors and to choose optimal refinement strategy. I observed that this often results in reducing the final model refinement to the improvement of PDB “sliders”, graphically combining several global model quality indicators, which may obscure real, local problems.

One of the most elusive errors in macromolecular models are register shifts, where the backbone is traced correctly, but residues are systematically assigned the identity of a residue a few amino acids up or down in sequence (Wlodawer *et al*., 2018). The issue can be easily detected in high resolution structures as it causes significant mismatches between the model and the electron-density map for several neighbouring sidechains. Moreover, prominent difference-density peaks indicate missing or excess side-chain atoms in the model. At lower-resolutions, however, deteriorated map-model fit due to wrongly modelled sidechains can be easily mistaken for poorly resolved model fragments. Difference-density peaks are usually weaker and visible only for a few well resolved sidechains. The effect on global model-data fit scores (R-free or R-work/R-free gap) can be detectable but is typically small as the number of excess or missing atoms is usually negligible compared to the overall size of a model. Register shifts resulting from a tracing error (deletion or insertion) can also be detected by the presence of backbone geometry outliers. For example, deletions are often compensated with a stretched backbone, which after refinement may result in a twisted peptide bond (in COOT, clearly marked with a yellow polygon). Similarly, wrongly assigned sidechains often result in severe steric clashes that cannot be corrected during refinement. In summary, register shifts often produce multiple model-validation metric outliers simultaneously, none of them unambiguous. Therefore, correct identification of the source of the problem usually requires a tedious, residue-by-residue analysis of a map and crystal structure model by an experienced crystallographer (Croll *et al*., 2021). In cases where lower resolution maps provide little help in model validation, the recently developed conkit-validate may an attractive option as it is based on purely geometrical comparison of model-derived and AI-predicted intramolecular contacts and distances (Sánchez Rodríguez *et al*., 2022).

In a recent publication I presented *checkMySequence*, a tool for the automated detection of register-shift errors in cryo-EM models (Chojnowski, 2022). The method is based on *findMySequence*, a protein sequence identification tool for crystallography and cryo-EM (Chojnowski *et al*., 2022). The *checkMySequence* algorithm systematically assigns input-model fragments to a reference sequence to identify regions where the new sequence-assignment challenges the sequence-assignment hypothesis in the input model. This approach provides a conceptually simple and fast tool for automated detection of register-shifts in cryo-EM models, including very large macromolecular complexes (e.g. complete ribosomes). Here, I show that the same approach can be applied to the analysis of MX models, using refined coordinates and standard, model-bias corrected combined crystallographic 2mFo-DFc maps. I describe in detail five crystal structure models deposited in the PDB with register shift errors that can be unambiguously detected using *checkMySequence* but would be difficult to identify automatically using other available model validation methods.

## 2. Materials and methods

### 2.1. Crystal structure benchmark set

For benchmarks, I selected from the PDB crystal structure models of proteins, with or without nucleic acid components, solved at a resolution between 2.0 and 3.0Å. I considered only models deposited with corresponding diffraction data. Out of 68,955 models fulfilling these criteria as of June 14th, 2022 I randomly selected 10,000 structures and downloaded from PDBe (Armstrong *et al*., 2020) corresponding atomic coordinates and amino-acid sequences in mmCIF and FASTA formats respectively. Maximum likelihood Fourier coefficients for combined (2mFo-DFc) and difference (mFo-DFc) maps calculated using REFMAC5 (Murshudov *et al*., 2011) and DCC (Yang *et al*., 2016) were downloaded in MTZ format from the RCSB (Burley *et al*., 2019).

### 2.2. Selection of test fragments

The performance of sequence assignment procedure implemented in *findMySequende* was tested using a large set of continuous, protein-chain test-fragment. From each of 13,525 unique protein chains in the benchmark set three continuous fragments were selected at random giving in total two sets of 40,575 test-fragments of 10 and 20 residues. Fragments with unknown residues, marked as ‘UNK’ in the model, were rejected. Fragments where the number of residues differed from the difference between flanking residue indices (possibly non-continuous) were rejected as well. This resulted in small differences between the expected and observed number of test-fragments.

### 2.3. Data analysis and processing software

Benchmark set structures were analysed fully automatically using *checkMySequence* version 1.4.1 and *findMySequence* version 1.0.8. Structural models with plausible sequence-register errors described in this work were analysed and rebuilt interactively using COOT version 0.9.8.4 and CCP4 8.0.005 within CCP4 Cloud version 1.7.006 (Krissinel *et al*., 2022). Unless otherwise stated, corrected models were refined automatically using REFMAC5 version 5.8.0267 and PDB_REDO version 7.38 (Joosten *et al*., 2014). Figures were prepared using PyMOL (DeLano, 2002) and matplotlib (Hunter, 2007). Structural superposition was performed using GESAMT version 1.18 (Krissinel, 2012) and corresponding root-mean-square deviation (RMSD) values were calculated using CA-atoms.

## 3. Results and discussion

### 3.1. Sequence assignment statistics

The *checkMySequence* program systematically aligns continuous fragments of an input protein model to the target sequence based on the corresponding map. The program internally uses an algorithm implemented in the *findMySequence* that scores each sequence alignment with a p-value – a probability that an alignment was observed by chance. Cases where the p-value is smaller than a predefined threshold and the new sequence alignment is different from the input model may indicate a register shift. I have recently shown (Chojnowski, 2022) that this approach can reliably identify register shift errors in cryo-EM models. Unlike cryo-EM, however, MX electron-density maps are calculated using phase information derived from atomic models. This inevitably results in a model-bias and the presence of the electron density map features derived from the model and not from the experimental data, which may obscure errors. Although the model-bias issue is addressed with the maximum likelihood maps, commonly used for MX model building and interpretation, it was not clear whether and to what extent it would affect the performance of the AI-based classifier implemented in the *findMySequence*. In particular, the p-value threshold determined for the analysis of cryo-EM models needed to be redefined for the MX models.

In the first step I analysed the distribution of p-values for test-fragments randomly selected from benchmark structures as described in Material and Methods section. The number of test-fragments for which re-assigned and model sequences differed was relatively small; 389 out of 39,774 and 197 out of 38,718 test-fragments of 10 and 20 residues respectively. The number of test-fragments with misassigned sequences is also significantly less than observed previously for EM structures (Chojnowski, 2022). This agrees with estimated accuracy of residue-type classifiers used in the *findMySequence*, which is noticeably higher for MX than cryo-EM (Chojnowski *et al*., 2022).

The test-fragments with correctly and wrongly assigned sequences are clearly separated by the p-value, which is an indicator of the strong predictive power of the classifier (Figure 1). For the sake of simplicity, the threshold defined previously for cryo-EM structures (p-value of 0.14) was also used for the MX structures. In the current MX structures benchmark set, this threshold corresponds to a 98.0% and 99.7% one-sided confidence interval for a correct sequence assignment for fragments of 10 and 20 residues respectively (Figure 1). Moreover, less than 0.1% of test-fragments were assigned a wrong sequence with p-value below this threshold (regardless of fragment length). Even though it is not known at this stage how many of these originate from structures with sequence register issues, they correspond to model fragments that are very ‘unusual’ in statistical terms and deserve closer attention.

**Figure 1.**
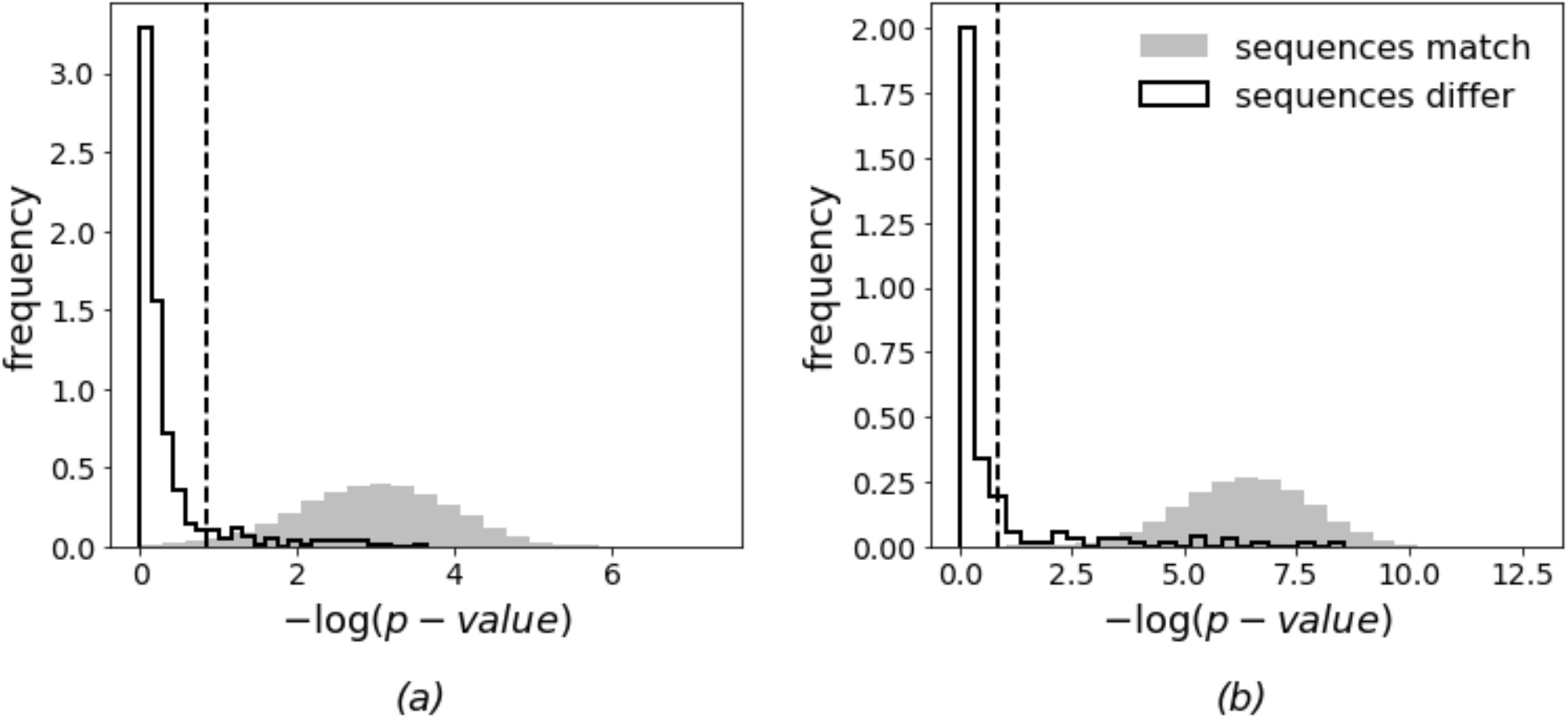
Statistics of sequence re-assignment of randomly selected continuous protein chain fragments from benchmark set MX structures. Grey and contoured histograms represent cases where newly assigned sequences match or differ respectively from the reference model for test-fragments of (a) 10 and (b) 20 amino acids. Vertical dashed line depicts a standard threshold used by *checkMySequence* for outlier identification in cryo-EM models. The plots’ ordinate axes show −log(p-value); higher values correspond to lower p-values and more reliable sequence assignments. Frequency histograms are shown for clarity, but the sets presented on each panel are strongly unbalanced. The number of test-fragments with re-assigned sequences that don’t match the reference model is 1% of the overall number of test fragments in the benchmark set.

### 3.2. Benchmark set analysis with *checkMySequence*

The *checkMySequence* was used to systematically scan all the crystal structures in the benchmark set, with the parameters derived in the previous section, deposited models, and corresponding maximum likelihood 2mFo-DFc maps. The analysis of the input structures took 18s on average and less than 105s for 99% of the tasks. Overall, the program identified sequence-assignment issues in 264 out of 10,000 structures from the benchmark set. They include 26 structures with residue indexing issues, such as an unmodelled loop ignored in residue numbering (no gap) and 86 structures with sequence mismatches or unidentified residues in a model. In 89 structures, at least one chain of 10 or more amino acids could not have been assigned to a reference sequence. Given the high sensitivity of *findMySequence*, which was used here to identify reference sequences, this may indicate chains that are very poorly resolved in the electron density (Chojnowski *et al*., 2022). Finally, *checkMySequence* identified plausible register shifts in 70 structures. Of these, I selected five models, in which I corrected register shift errors using interactive modelling software. They are presented in detail below.

#### Case study 1: WD40-repeat domain from *Thermomonospora curvata*

The WD40-repeats are a large family of proteins with a variable-size beta propeller fold. The WD40-repeat protein from *Thermomonospora curvata* contains seven blades. The deposited crystal structure model (PDB entry 5yzv, (Shen *et al*., 2018)) with five molecules in the Asymmetric Unit (ASU) was solved by Molecular Replacement (MR) with Phaser program (McCoy *et al*., 2007) and refined at 2.5Å resolution with clearly elevated validation scores (*clashscore* of 27 and R/R-free values of 0.225/0.259). An automated processing with PDB_REDO didn’t improve the validation scores (*clashscore* of 36 and R/R-free values of 0.199/0.259), indicating that the refinement strategy wasn’t an issue here. The different molecules in the ASU have also a relatively large structural variability reaching 1.2Å RMSD, which is unexpected given the compact fold of the crystallised protein.

The *checkMySequence* analysis of the deposited coordinates revealed multiple register shifts in 3 out of 5 molecules in the ASU. The alternative sequences were assigned with a p-value below 0.01 and are therefore of high confidence (Figure 1). However, the suggested register shifts were unusually large (up to 200 residues) and inconsistent between neighbouring chain fragments, even though no clear tracing issues were visible in the model. A more detailed inspection of the model and map revealed a number of sidechains with strong difference-density peaks (e.g. Trp A/503 Figure 2b), confirming that the structure may indeed suffer from an unusual modelling issue.

**Figure 2.**
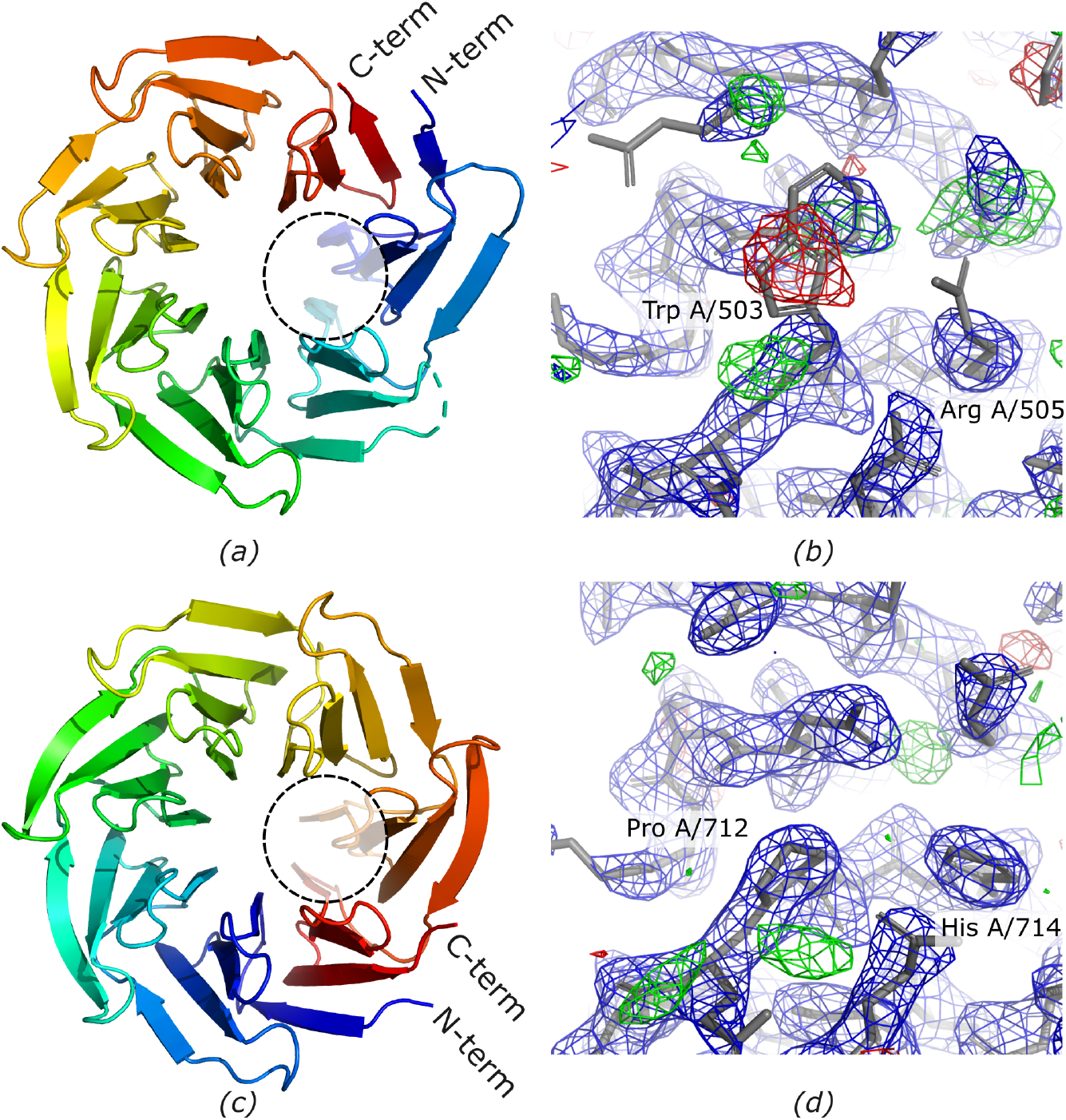
Comparison of deposited (a,b) and corrected (c,d) models of WD40-repeat domain from *T. curvata*. Three molecules in ASU of the deposited crystal structure were rotated about a seven-fold pseudo-symmetry axis of the structure (a,c) resulting in a number of clear density outliers in the model e.g. Tryptophan A/503 and Arginine A/505 labelled on panel (b). Correcting the molecule rotation results in a much better fit to the data (d). Model fragments shown on panels (b) and (d) are indicated by dashed circles on panels (a) and (c) respectively. The combined 2mFo-DFc (blue) and difference mFo-DFc (red/green) maximum likelihood maps calculated using REFMAC5 are shown at 1.5σ and 3σ levels respectively.

The source of the problem turned out to be a rotation about a seven-fold pseudo-symmetry axis of the protein resulting in an inconsistent register of beta-propeller blades in the model (Figure 2a,c). The sequence differences between WD40-repeat blades were presumably obscured by the presence of an approximate seven-fold symmetry of the backbone during the MR search. This resulted in a partially incorrect sequence register of three molecules in the initial MR solution that was overlooked during the subsequent refinement steps. To confirm this, I corrected the model following suggestions from the *checkMySequence* analysis. As the refinement of register-shifted chains resulted in their deformation (chains A, C, and D have RMSD over 1.0Å when compared to chains with a correct register) I replaced them with a very reliable prediction from AlphaFold Protein Structure Database (release v3 for UniProt entry P49695 with pLDDT over 90; (Varadi *et al*., 2022)). To enforce a correct sequence register I superposed the prediction onto the chains after re-assigning deposited model chains to a target sequence using *findMySequence*. Refinement of the corrected model using REFMAC5 with jelly-body restraints required 150 cycles to converge but resulted in a notably better-quality scores compared to deposited coordinates; R/R-free values reduced to 0.170/0.204 (from 0.225/0.259) and *clashscore* to 4 (from 27). The refinement resulted in a relatively small changes in the chains replaced with the initial AlphaFold2 model prediction (0.4Å RMSD compared to the initial model). The overall quality of the map-model fit, however, improved noticeably (Figure 2d). In contrast to the deposited coordinates, different WD40-repeat molecules in the final crystal structure model are also virtually identical with RMSD differences not exceeding 0.45Å, which shows that the initially observed structural diversity was indeed a consequence of sequence misassignment.

#### Case study 2: *Helicobacter pylori* helicase with degraded helices

The structure of DnaB helicase from *Helicobacter pylori* (*Hp*DnaB) consists of two globular domains separated by a linker forming helices 7, 8, and 9. The deposited crystal structure model (PDB entry 3gxv, (Kashav *et al*., 2009)) with two molecules in the ASU was solved using MR using Phaser program and the N-terminal domain of a related helicase from *Mycobacterium tuberculosis* as a search model (PDB entry 2r5u, 25% sequence identify). The final model was refined at 2.5Å resolution to reported R/R-free of 0.249/0.278 with *clashscore* of 32. An optimalization of refinement strategy with PDB_REDO reduced *clashscore* to 4.57 at the expense of slightly worse R/R-free factors of 0.262/0.291.

The crystallised *Hp*DnaB variant consists of a globular N-terminal domain (NTD) and helix 7. The NTD and helix 7 dimer in the ASU is stabilised by two short, helical peptides, which the structure authors identified as the helix 7 degraded from a complete construct. This was further confirmed by crystal electrophoresis and mass spectrometry experiments. It is worth noting that alternatively to author’s interpretation the crystal content can be inhomogeneous, having in the ASU a mixture of degraded helices 7 and additional complete *Hp*DnaB molecules with disordered NTD domains. This seems plausible given the crystal packing, the very high solvent-content of the deposited crystal structure (72.4%) and results of Matthews coefficient analysis, which suggests four HpDnaB molecules (NTD and helix 7) in the ASU. This would explain relatively high R/R-free factors of the deposited model.

One of the two isolated helices 7 in the crystal structure (chain C) is a very prominent register-shift in the *checkMySequence* analysis, with the alternative sequence assignment p-value below 0.001 (Figure 1). This is confirmed by the presence of strong difference density peaks suggesting that several sidechains are misaligned in the model, for example a Phenylalanine C/111 and Asparagine C/115 (Figure 3a).

**Figure 3.**
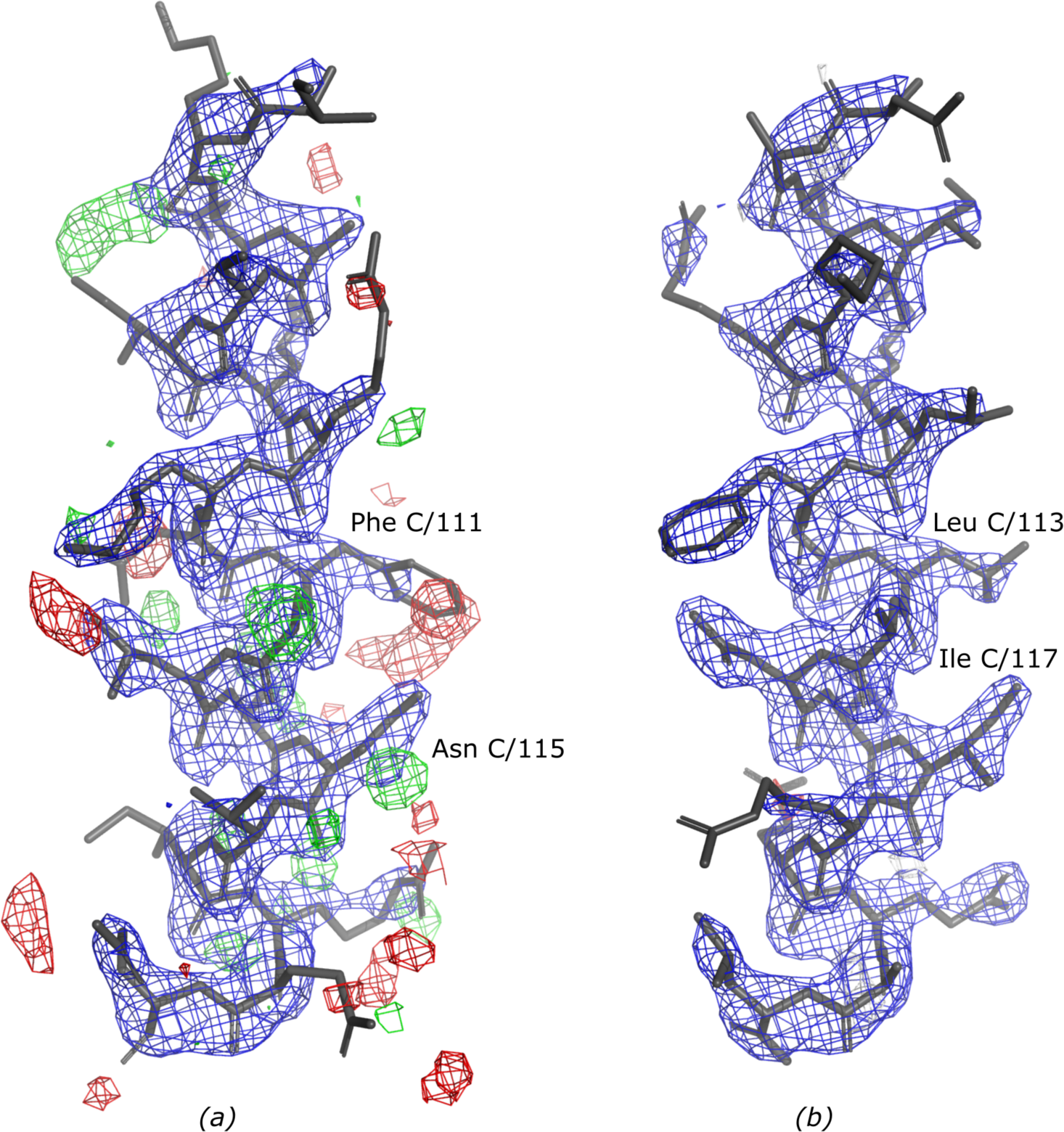
Crystal structure of an isolated helix 7 from the N-terminal domain (NTD) of *H. pylori* DnaB (*Hp*DnaB) helicase. Comparison of deposited (a) and corrected (b) models. The peaks in the difference density mFo-DFc map can be eliminated by shifting the model sequence register by 2 residues. For example, clearly too large Phenylalanine C/111 resulting in a prominent negative (red) difference density peak is replaced with a smaller Leucine C/113 in the model with corrected sequence register. Similarly, a clear positive peak (green) near Asparagine C/115 is interpreted with an Isoleucine C/117 sidechain of the corrected model. The combined 2mFo-DFc (blue) and difference mFo-DFc (red/green) maximum likelihood maps calculated using REFMAC5 and PDB_RDO are shown at 2σ and 3σ levels respectively. At this threshold no difference density map features are visible in the presented ASU fragment of the corrected model.

Automated refinement with PDB_REDO of the model with helix 7 (chain C) re-assigned to the target sequence with *findMySequence* resulted in a much better fit of the coordinates to the corresponding 2mFo-DFc map and in a reduction of strong difference density peaks (Figure 3b). This suggests that the new sequence register indeed fits better the data. Surprisingly, the correction of model sequence resulted in a negligible reduction of validation scores (R/R-free factors of 0.257/0.289 and *clashscore* of 7.55). This, however, can be attributed to a relatively small (albeit important for model interpretation) modification of the coordinates (difference of 18 out of over two thousand non-H atoms) and relatively high overall R-values as discussed above.

#### Case study 3: A hydrogenase from *Thermosipho melanesiensis*

The HydF is one of maturation proteins required for an activation of a [FeFe] hydrogenase (HydA). The structure of *Thermosipho melanesiensis* HydF (TmeHydF) is composed of three domains (Figure 4a); dimerization (residues 7–166), GTP-binding (residues 172–262) and cluster-binding domain (residues 263–395). The structure of the protein in complex with an Fe-S cluster (PDB entry 5kh0, (Caserta *et al*., 2017)) was solved by MR using a closely related homologue, an apo-HydF structure from *T. neapolitana* as a search model (PDB entry 3qq5; 97% sequence identity). The deposited model with four molecules in the ASU was refined at 2.8Å resolution to reported values of R/R-free and *clashscore* of 0.233/0.262 and 5.55 respectively. Automated refinement of the deposited model with PDB_REDO resulted in R/R-free of 0.225/0.261 and *clashcore* 17.53.

**Figure 4.**
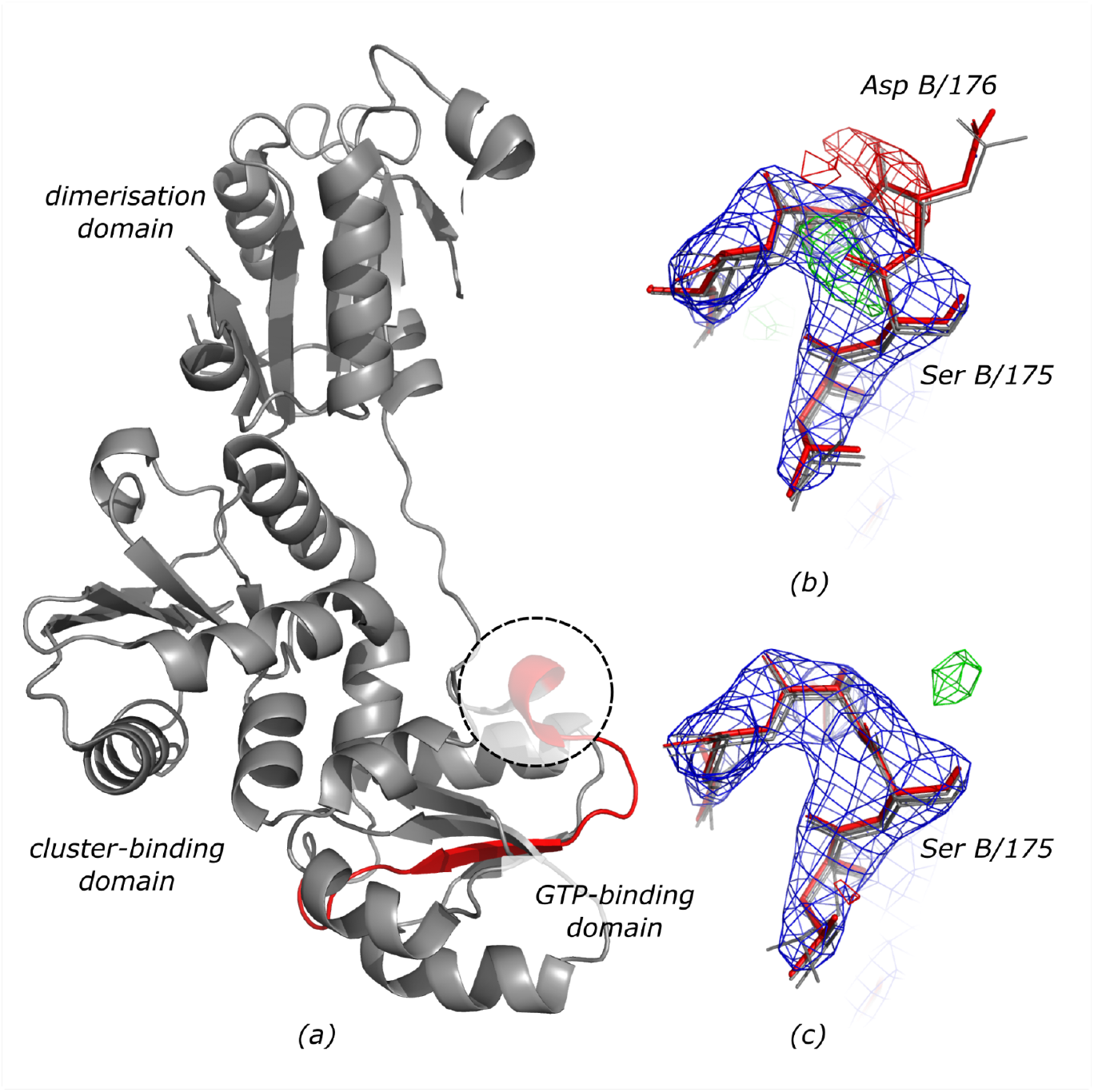
Crystal structure of HydF maturase with a register-shifted fragment of a GTP-binding domain shown in red (a). The dashed circle indicates a region of the deposited model shown on panel (b) with corresponding electron-density maps. The Aspartate B/176 is a clear map-model fit outlier that results in a strong negative peak in difference-density map and a register shift in a fragment shown in red in panel (a). The remaining three HydF chains in the ASU, superimposed onto chain B are shown in grey. After correcting the model and subsequent restrained refinement the map-model agreement clearly improves (c). The combined 2mFo-DFc (blue) and difference mFo-DFc (red/green) maximum likelihood maps calculated using REFMAC5 and PDB_REDO are shown at 1.5σ and 3σ levels respectively.

An analysis with *checkMySequence* identified plausible register shifts with a p-value below 0.001 in a beta-strand between residues 178 and 194 at N-terminal of the GTP-binding domain in all four TmeHydF chains in the ASU (Figure 4a). Closer inspection revealed that the shift is caused by a clear insertion at a residue 176 in chains B and D (Figure 4b) or a mistraced region between residues 180 and 182 in chains A and C (not shown). The insertions are further compensated by deletions at the same position in all four chains (residue 192). The register-shifted region doesn’t affect the conformation of the protein’s active site in the cluster-binding domain, which was analysed in more detail by the authors of the structure. After correcting the register-shifted fragments with *findMySequence* and COOT, and subsequent restrained refinement in PDB_REDO with REFMAC5, the R/R-free improved slightly to 0.221/0.252 (from 0.225/0.261) and the *clashscore* reduced to 11 from 17. Moreover, the clearly visible, prominent difference-density peaks in regions corresponding to wrongly assigned sidechains disappeared (Figure 4c).

The TmeHydF structure prediction downloaded from AlphaFold Protein Structure Database (release v3 for UniProt entry A6LMQ7) has a different orientation of the dimerization domain relative to the remaining two domains to the extent that would make the use of the AlphaFold2 prediction for model building or as a MR search model not straightforward. Interestingly, this is not reflected in the Predicted Alignment Error (PAE) plot, which would additionally complicate the model usage. The predicted and corrected crystal structure models, however, agree very well locally. For example, the GTP-binding domains superpose with RMSD of 0.78Å with the only significant differences in a loop following the register-shifted region (predicted with a low accuracy; pLDDT below 50). Thus, the prediction could in principle be used to identify and correct the register shift error in the deposited model.

#### Case study 4: Protein L31e from large ribosomal subunit of *Haloarcula marismortui*

The crystal structure of the 50S large ribosomal subunit of *Haloarcula marismortui* has been determined at 2.65Å resolution and refined to R/R-free of 0.176/0.214 and *clashscore* of 16 (PDB entry 1yi2, (Tu *et al*., 2005)). An analysis with *checkMySequence* revealed that a C-terminal fragment of a peripheral ribosomal protein L31e may be shifted by 2 residues (p-value 0.1) between residues X/77 and X/88 (the last modelled residue in the chain). A closer inspection of the deposited model and maps revealed several difference-density peaks in the C-terminal fragment of the protein (Figure 5a). After re–assigning the fragment to the target sequence with the *findMySequence*, rebuilding a short loop preceding it in COOT, and subsequent refinement with PDB_REDO the overall map-model agreement of the chain clearly improved (Figure 5b). The final model R/R-free didn’t change compared to values obtained using PDB_REDO for the deposited coordinates (0.166/0.206 versus 0.167/0.206), *clashscore* reduced slightly from 4.33 to 3.29.

**Figure 5.**
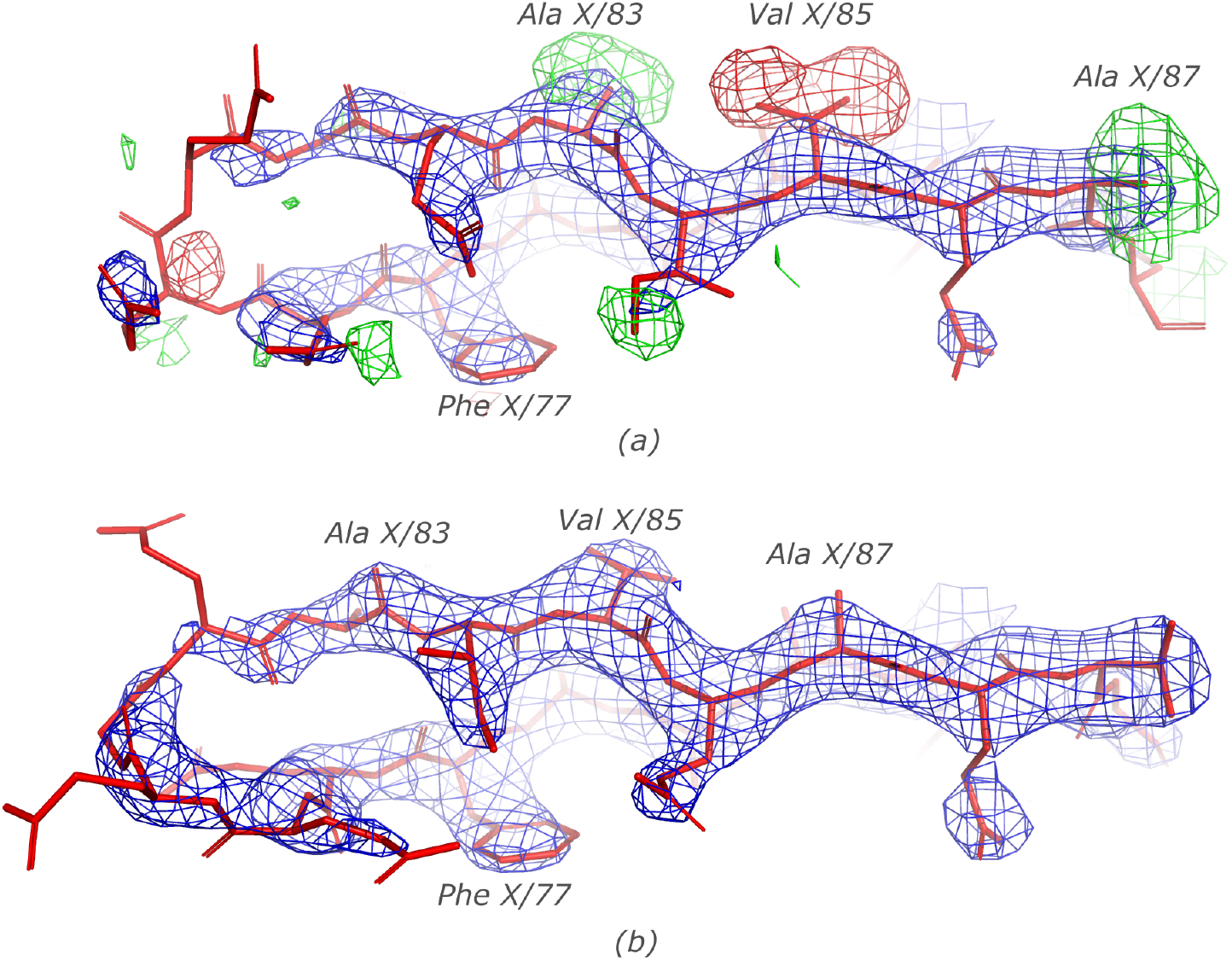
Ribosomal protein L31e from a crystal structure of a large ribosomal subunit from *H. marismortui*. A solvent-exposed, poorly resolved loop following Phenylalanine X/77 was traced too short in the original model that resulted in a 2-residue sequence register shift in a C-terminal part of the chain. Strong difference density peaks showing a few excess and missing atoms in the deposited structure for Alanine X/83, Valine X/85, and Alanine X/87 (a) disappear after refining the model with corrected sequence register (b). The combined 2mFo-DFc (blue) and difference mFo-DFc (red/green) maximum likelihood maps calculated using REFMAC5 and PDB_REDO are shown at 1.5σ and 3σ levels respectively.

A structure prediction from AlphaFold2 database (release v3 for UniProt entry P18138) has overall high confidence and agrees very well with the corrected crystal structure model (RMSD 0.47Å including the poorly resolved loop that had to be rebuilt). This region, however, was scored slightly lower than the remaining structure (pLDDT between 80 and 85).

#### Case study 5: A glutaminase from *Geobacillus kaustophilus*

The structure of a glutaminase from *Geobacillus kaustophilus* was refined at 2.1Å resolution with four molecules in the ASU (PDB entry 2pby) to R/R-free of 0.195/0.249 and *clashscore* 7.62, which reduced to 0.174/0.207 and *clashscore* 6.44 after automated refinement strategy optimization with PDB_REDO. The *checkMySequence* analysis revealed an unambiguous shift of sequence register in chain B between residues B/54 and B/92 (p-value less than 0.0001). The model inspection revealed an insertion at a residue B/57 starting a register-shift continuing till a chain break at a residue B/92 (Figure 6a and 6b). As the structure was solved by a Structural Genomics Initiative (RSGI/SECSG) and remains unpublished little is known about structure determination details. According to the PDB file header it was solved by MR using EPMR (Kissinger *et al*., 2001) and the structure of a related glutaminase from *Bacillus subtilis* as a search model (PDB entry 1mki). The two structure models are very similar and 275 out of 291 and 321 residues in target and search models respectively align with 1.1Å RMSD. Nevertheless, they share only 46% sequence identity and have multiple loops of different length and/or conformation, which suggests that the deposited model underwent an extensive (possibly automated) rebuilding that may have resulted in the register shift. All four chains in the model are virtually identical (0.23Å RMSD) with the only visible difference in a conformation of residues flanking a solvent exposed, disordered loop between residues 92 and 109, and the clearly visible deletion in chain B that resulted in a register shift (Figure 6a).

**Figure 6.**
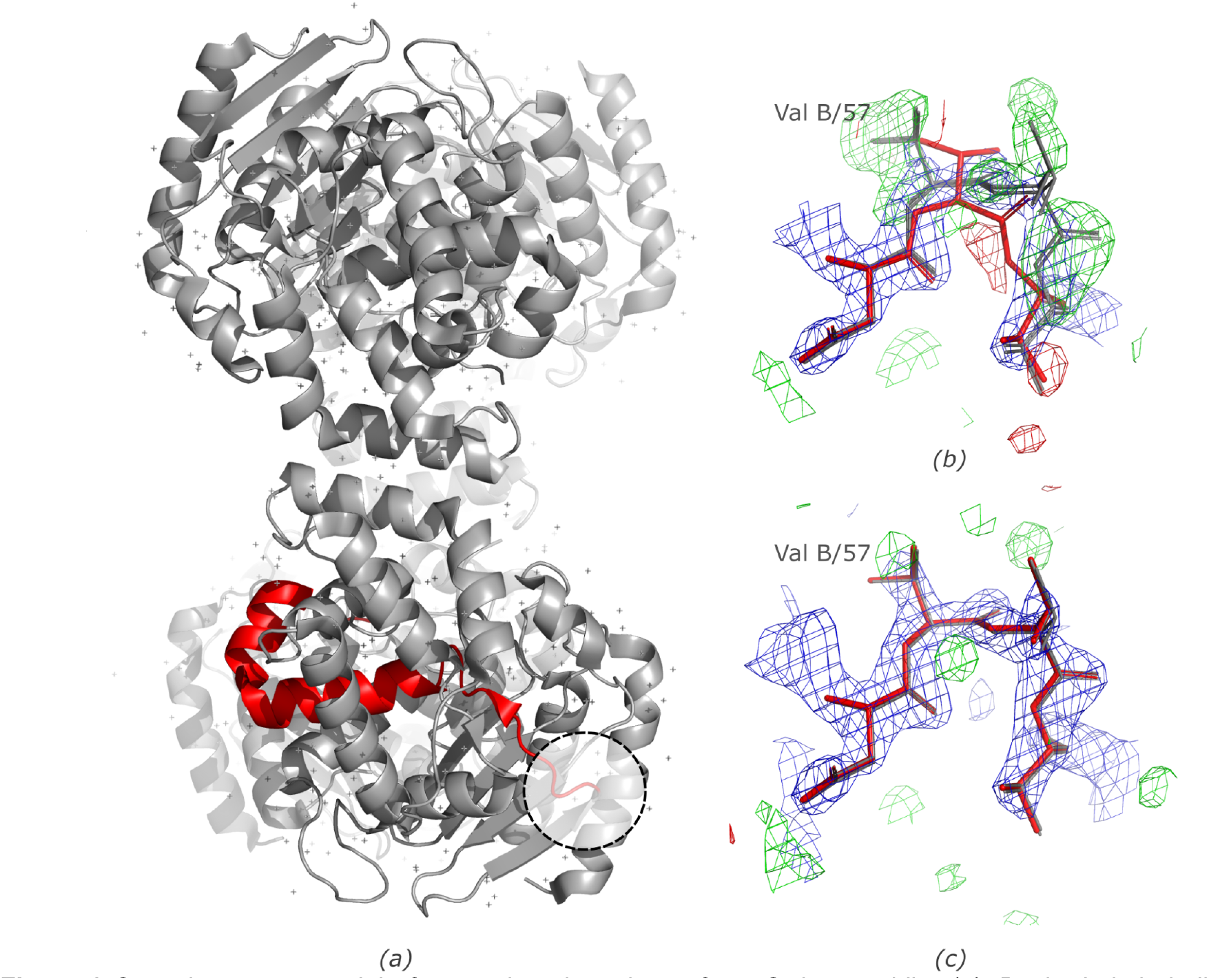
Crystal structure model of a putative glutaminase from *G. kaustophilus* (a). Dashed circle indicates a region of the deposited model shown on panel (b) with corresponding maps. A deletion near Valine B/57 results in multiple strong positive difference-density map peaks and a register shift in a fragment shown in red in panel (a). The panel depicts residues 56-59 from chain B (red) and superposed models of remaining chains A, C, and D (grey). After correcting the model and subsequent restrained refinement the map-model agreement clearly improves (c). The combined 2mFo-DFc (blue) and difference mFo-DFc (red/green) maximum likelihood maps calculated using REFMAC5 and PDB_REDO are shown at 2σ and 3σ levels respectively.

After correcting the main-chain tracing issue in chain B, re-assigning the register-shifted model fragment to the sequence using *findMySequence* and subsequent automated refinement using REFMAC5 and PDB_REDO the R/R-free and *clashscore* reduced to 0.168/0.197 and 1.90 respectively (from 0.174/0.207 and 6.44 after initial PDB_REDO optimization). In addition, a better map-model fit was obtained, as well as a better agreement between all four molecules in the ASU (Figure 6c).

## 4. Conclusions

The purpose of building scientific models is to enable the interpretation of complex experimental data in the light of the available theoretical knowledge. Consequently, models are always provisional and can be updated if new evidence becomes available. This applies to the structural models of macromolecules; they are tentative and can be always improved; with better data, by a laborious iterative refinement or with a new, more robust data analysis software.

Here, I presented *checkMySequence*– a fast and fully automated method for the identification of register shifts in crystal structure models of proteins. I showed that it can identify errors in structural models that were already considered “good enough” and deposited to the PDB. The sequence assignment issues I selected to describe in detail do not affect the conclusions derived from corresponding models by their authors as they were found in peripheral regions (ribosomal protein L31e), affect overall model quality (WD40-repeat), or only one out of multiple protein copies in the ASU (glutaminase). It is probably for this reason that they went unnoticed in the first place. It is not clear, however, how these errors affected or will affect subsequent studies. It also remains to be seen how many models deposited in the PDB have register errors that affected their functional analysis. This cannot be studied *en masse* as it requires an individual approach by specialists, either revisiting their own models, or aggregating available structural data. For others a simple warning about a potential sequence-assignment issue in a PDB deposited structure can help to avoid problems. I believe that *checkMySequence* will prove helpful to all of them.

Although all the described issues could have been deduced from the presence of prominent difference density peaks, unusual backbone geometry, local differences between different copies of the same molecule in ASU, high *clashscore* or elevated R-factors, only *checkMySequence* clearly annotated the errors. This should make *checkMysequence* particularly useful for inexperienced users or when validated models are very large, like the 50S ribosomal subunit presented above, where a detailed residue-by-residue analysis of map-model fit and model geometry is not feasible.

I’ve also shown that a few of the presented errors can be corrected (and possibly could have been avoided) using AI-based predictions for the structure determination, as has already become a standard. In some cases, however, this would not be enough to avoid an error. For example, when an isolated fragment needs to be assigned to target sequence (the case of HpDnaB), or an error is obscured by an unusual fold of a protein (WD40-repeat). In such cases *checkMySequence* will prove especially useful.

The presented results were restricted to crystal structures determined between 2.0 and 3.0Å resolution, which dominates among MX structures deposited in the PDB. In this resolution range all map-model fit problems are usually clearly visible, although not always trivial to identify and correct. It is also relatively easy to present visual evidence that the new model does indeed better explain the experimental data. At lower resolutions this becomes increasingly difficult. Therefore, a more challenging analysis of register errors in low resolution crystal structures, probably far more frequent, I leave for future collaborative work and a more robust methodology combining an approach presented here and an orthogonal method restricted to model-geometry analysis (Sánchez Rodríguez *et al*., 2022).

## 5. Data and code availability

The latest version of *checkMySequence* source code and installation instructions are available at https://gitlab.com/gchojnowski/checkmysequence.

## 6. Acknowledgements

I would like to thank Katherine S. H. Beckham, Isabel Bento, and Daniel J. Rigden for critical reading of the manuscript and very helpful comments.

